# Host Chitinase 3-like-1 is a Universal Therapeutic Target for SARS-CoV-2 Viral Variants in COVID 19

**DOI:** 10.1101/2022.01.21.477274

**Authors:** Suchitra Kamle, Bing Ma, Chang Min Lee, Gail Schor, Yang Zhou, Chun Geun Lee, Jack A. Elias

**Author notes:** **Author Contributions** Conception and design: SK, JAE, CGL; Generation of experimental resources and data collection: SK, BM, CML, GS, YZ; Analysis and interpretation: SK, CGL, JAE; Drafting the manuscript for important intellectual content: JAE, CGL, SK. **Competing Interests Disclosures:** Jack A. Elias is a cofounder of Elkurt Pharmaceuticals and Ocean Biomedical which develop therapeutics based on the 18 glycosyl hydrolase gene family.

## Abstract

COVID 19 is the disease caused by severe acute respiratory syndrome coronavirus-2 (SARS-CoV-2; SC2) which has caused a world-wide pandemic with striking morbidity and mortality. Evaluation of SC2 strains demonstrated impressive genetic variability and many of these viral variants are now defined as variants of concern (VOC) that cause enhanced transmissibility, decreased susceptibility to antibody neutralization or therapeutics and or the ability to induce severe disease. Currently, the delta (δ) and omicron (o) variants are particularly problematic based on their impressive and unprecedented transmissibility and ability to cause break through infections. The delta variant also accumulates at high concentrations in host tissues and has caused waves of lethal disease. Because studies from our laboratory have demonstrated that chitinase 3-like-1 (CHI3L1) stimulates ACE2 and Spike (S) priming proteases that mediate SC2 infection, studies were undertaken to determine if interventions that target CHI3L1 are effective inhibitors of SC2 viral variant infection. Here we demonstrate that CHI3L1 augments epithelial cell infection by pseudoviruses that express the alpha, beta, gamma, delta or omicron S proteins and that the CHI3L1 inhibitors anti-CHI3L1 and kasugamycin inhibit epithelial cell infection by these VOC pseudovirus moieties. Thus, CHI3L1 is a universal, VOC-independent therapeutic target in COVID 19.

## INTRODUCTION

Coronavirus disease 2019 (COVID 19), the illness caused by severe acute respiratory syndrome coronavirus virus-2 (SARS-CoV-2; SC2), was first discovered in man in 2019 and declared a global pandemic by the World Health Organization (WHO) on March 11, 2020. It is the cause of a global health crisis with countries experiencing multiple waves of illness resulting in more than 273 million confirmed clinical cases and more than 5.34 million deaths as of December 17, 2021 (1-8). The disease caused by SC2 was initially noted to manifest as a pneumonia (9). It is now known to have impressive extrapulmonary manifestations and vary in severity from asymptomatic to mildly symptomatic to severe disease with organ failure to death (10, 11). However, the cellular and molecular events that account for the multiple waves of disease and the impressive clinical and pathologic heterogeneity that have been seen have not been defined.

SC2 interacts with cells via its spike (S) protein which binds to its cellular receptor angiotensin converting enzyme 2 (ACE2) (12-15). To mediate viral entry the S protein is processed into S1 and S2 subunits by the S priming proteases (SPP) including TMPRSS2, cathepsin L (CTSL) and to a lesser degree, FURIN. The S2 subunit mediates the fusion of the viral envelope and cell plasma membrane to allow for virus-cell entry (14, 16, 17). In keeping with the importance of virus-cell interactions, many treatments for COVID 19 have focused on disease prevention using non-pharmacologic public health measures, antiviral antibodies and a critical global vaccination strategy (8). Treatments of acute infection include supportive interventions, anti-inflammatories, recently described oral antivirals and direct antivirals such as remdesivir (8). Surprisingly, although ACE2 and SPP play critical roles in SC2 infection and proliferation, therapeutics that focus on these host moieties have not been adequately investigated.

Early strains of SC2 from Wuhan China manifest limited genetic diversity (18). However, genetic epidemiologic evidence in February 2020 demonstrated the global emergence of a new dominant SC2 variant called D614G (18, 19). This variant was associated with enhanced transmissibility based on a S protein that is more likely to assume an “open” configuration and bind ACE2 with enhanced avidity when compared to the ancestral strain (8, 18). In the interval, since then multiple other SC2 variants have been appreciated. Many are now defined as variants of concern (VOC) due to their enhanced transmissibility, decreased susceptibility to neutralization by antibodies obtained from natural infection or vaccination, ability to evade detection or ability to decrease therapeutic or vaccine effectiveness (8). As of June 2021, 4 variants (alpha, beta, gamma and delta) had been defined as VOC. Most recently omicron has been added to the list of VOC. Of these viral moieties, delta and omicron (B.1.1.529) are most problematic. The delta variant has caused the deadly second wave of disease in India and waves of COVID 19 at other sites around the world (20, 21). It is also known to have a high level of transmissibility and virulence when compared to ancestral controls and the alpha (α), beta (β), and gamma (γ) variants. It manifests an enhanced ability to replicate and accumulates at very high levels in airways and tissues (8). In keeping with these characteristics, recent studies have also demonstrated that delta is associated with breakthrough infections in vaccinated individuals and a decrease in vaccine effectiveness, especially in the elderly (22). The omicron variant was first detected in November, 2021 and quickly declared a VOC based on its impressive transmissibility (23). It has 37 S protein mutations in its predominant haplotype, 15 of which are in its receptor binding domain (RBD) which is the major target of neutralizing antibodies (23-25). Although it appears to cause less severe disease than delta, its impressive ability to spread and resist antibody neutralization has resulted in surges that run the risk of overwhelming health care systems worldwide. In spite of the important differences in the S proteins of the variants and the impressive importance of S and ACE2 in COVID 19, therapies that focus on host targets such as CHI3L1, ACE2 and SPP that are effective in multiple SC2 variants have not been adequately defined.

We hypothesized that therapies that target the host factors involved in SC2 infection like CHI3L1 can contribute to the control of COVID 19 induced by all viral variants that use ACE2. To test this hypothesis, we employed pseudoviruses that expressed S proteins from the α, β, γ, δ and o variants and assessed the ability of CHI3L1-based interventions to modify their ability to infect human lung epithelial cells. These studies demonstrate that CHI3L1 augments the expression and accumulation of ACE2 and SPP and augments epithelial infection by the α, β, γ, δ and o pseudovirus variants. They also demonstrate that anti-CHI3L1 and the small molecule CHI3L1 inhibitor kasugamycin both inhibit the expression and accumulation of epithelial ACE2 and SPP and, in turn, inhibit epithelial infection by pseudoviruses that contain the α, β, γ, δ and o S proteins.

## RESULTS

### CHI3L1 stimulates epithelial cell uptake of the G614 and the alpha, beta, and gamma pseudotyped SC2 viruses

Previous studies from our laboratory demonstrated that CHI3L1 is a potent stimulator of epithelial expression of ACE2 and SPP and epithelial cell viral uptake (26). To determine if the major S variants altered these responses, we compared the uptake of pseudovirus with ancestral and mutated S proteins by untreated and CHI3L1-treated Calu-3 cells. As can be seen in Figure 1A, CHI3L1 was a potent stimulator of the uptake of pseudovirus with the ancestral G614 S protein. Similar increases in Calu-3 cell pseudovirus uptake were seen when the S proteins that are characteristic of the α, β, or γ variants were employed (Figure 1, B-D). When viewed in combination, these studies demonstrate that CHI3L1 augments SC2 pseudovirus uptake when the ancestral D614G and α, β, or γ S protein mutations are present.

**Figure 1.**
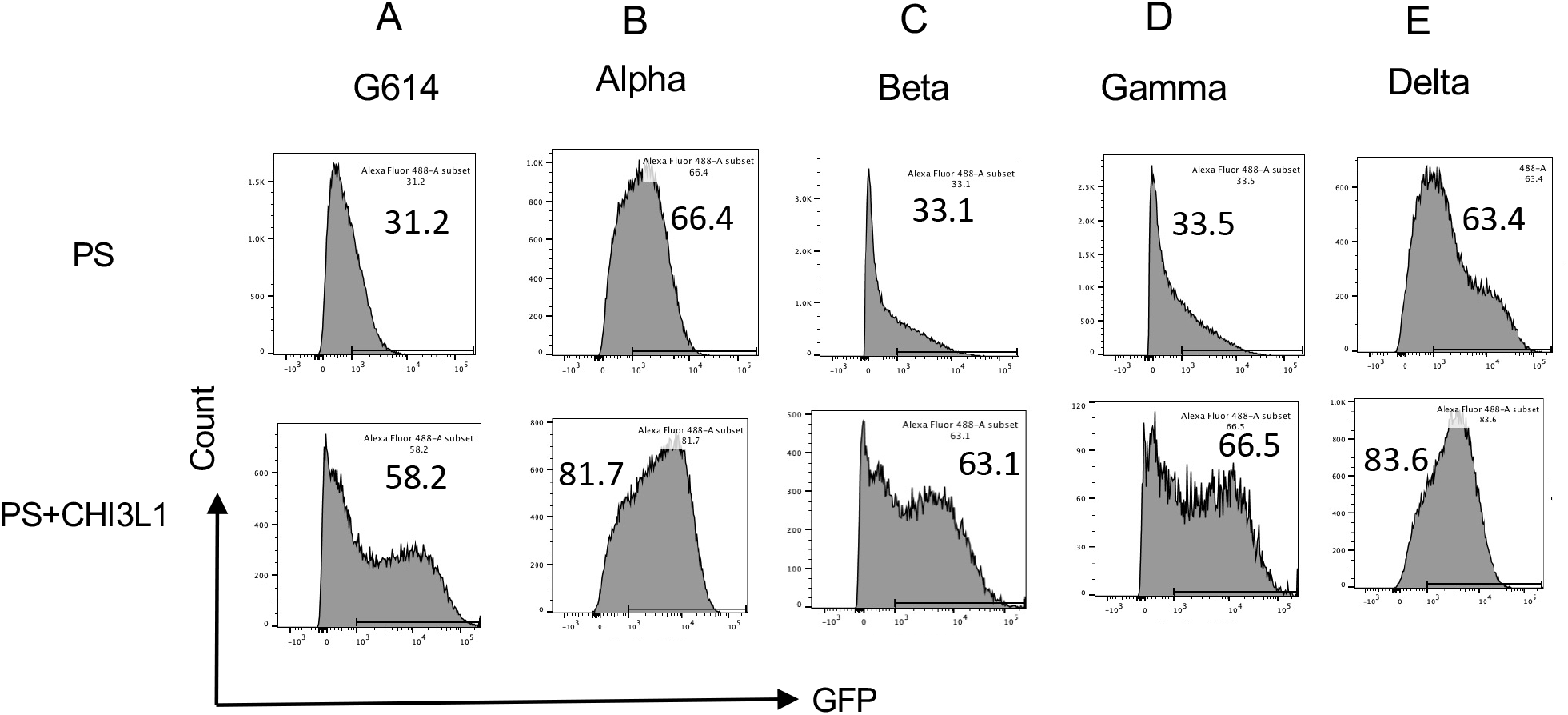
CHI3L1 stimulation of pseudovirus uptake. Calu-3 cells were incubated with recombinant human (rh) CHI3L1 (CHI3L1, 250ng/ml) or vehicle (PBS) control for 48 hours. Pseudoviruses (PS) that contain S proteins with G614, alpha, beta, gamma or delta mutations were added and GFP was quantitated by FACS. The % of GFP-positive cells was evaluated by flow cytometry. The noted values are representative of a minimum of 3 similar evaluations.

### CHI3L1 stimulates epithelial cell uptake of delta pseudotyped SC2 viruses

Because the delta SC2 variant manifests enhanced viral infectivity and has spread widely since it first appeared in December 2020, the ability of CHI3L1 to alter its ability to infect human epithelial cells was also assessed. In these experiments, CHI3L1 was also a potent stimulator of the uptake of pseudovirus with delta S protein mutations (Figure 1E). The findings noted above and these observations, in combination, demonstrate that CHI3L1 is a stimulator of human epithelial cell uptake of SC2 viral pseudotypes with S protein mutations from the α, β, γ and δ VOC.

### The monoclonal antibody “FRG” abrogates the CHI3L1-induced increase in epithelial cell uptake of the G614 and the alpha, beta and gamma pseudotyped viral variants

Studies were next undertaken to define the effects of the monoclonal anti-CHI3L1 antibody entitled “FRG” on the uptake of pseudovirus by Calu-3 cells treated with and without CHI3L1. As was seen with the ancestral G614 S protein mutation, treatment of Calu-3 cells with rCHI3L1 augmented pseudovirus uptake and FRG abrogated this increase while treatment with the IgG control did not (Figure 2A). Interestingly, FRG also diminished pseudovirus uptake by Calu-3 cells even when exogenous rCHI3L1 was not administered (Figure 2A). rCHI3L1 had similar stimulatory effects in experiments using pseudovirus with α, β, or γ S protein mutations (Figure 2, B-D). Importantly, the uptake of pseudoviruses with each of the S mutations in cells treated with and without rCHI3L1 was markedly diminished by FRG as well (Figure 2, A-D). When viewed in combination, these studies demonstrate that monoclonal anti-Chi3l1 targeting exogenous and or endogenous CHI3L1 effectively inhibits the uptake of pseudovirus with ancestral, α, β, or γ S protein mutations.

**Figure 2.**
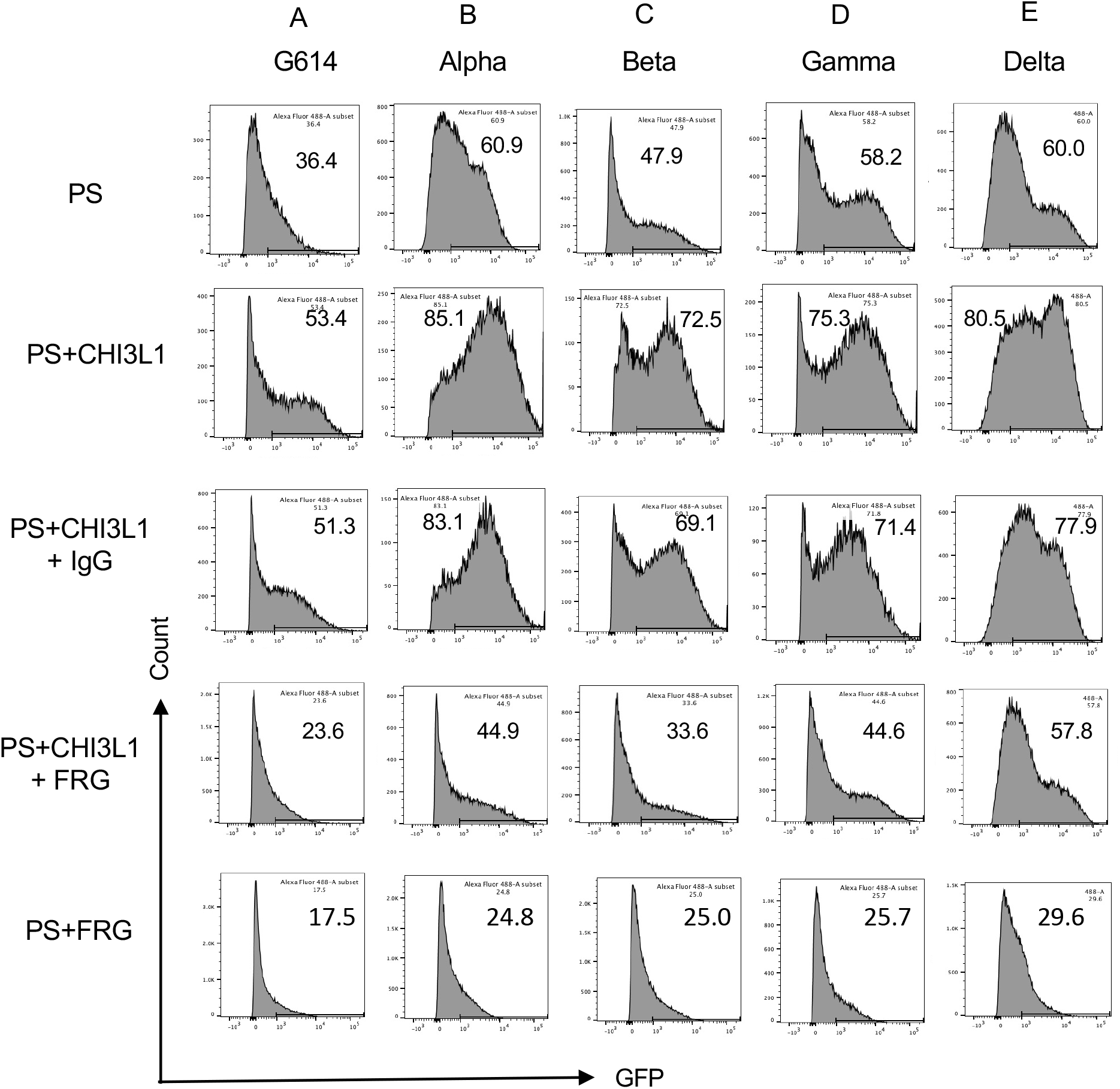
Effects of FRG on G614, alpha, beta, gamma and delta pseudovirus infection. Calu-3 cells were incubated with rhCHI3L1 (CHI3L1, 250ng/ml) or vehicle control for 48 hours in the presence of anti-CHI3L1 (FRG) or its isotype control (IgG). Pseudoviruses (PS) that contain S proteins with the G614, alpha, beta, gamma or delta mutations were added and GFP was quantitated by FACS. The % of GFP-positive cells was evaluated by flow cytometry. The noted values are representative of a minimum of 3 similar evaluations.

### The monoclonal antibody “FRG” abrogates the CHI3L1-induced increase in epithelial cell uptake of the delta pseudotyped viral variants

Because the delta SC2 variant has had such impressive clinical effects, the ability of FRG to alter its ability to infect human epithelial cells was also assessed. FACS-based evaluations demonstrated that CHI3L1 was a potent stimulator of the uptake of the pseudovirus with delta S protein mutations (Figure 2E). FRG abrogated this increase while treatment with the IgG control did not (Figure 2E). Interestingly, FRG also diminished pseudovirus uptake by Calu-3 cells even when exogenous CHI3L1 was not administered (Figure 2E). These findings were reinforced by immunocyotchemical evaluations. These studies demonstrated that CHI3L1 augmented Calu-3 cell ACE2 accumulation and delta pseudovirus infection (Figure 3). They also demonstrated that FRG abrogated the expression of ACE2 and delta pseudovirus infection at baseline and or after the administration of rCHI3L1 (Figure 3). When viewed in combination, these studies demonstrate that monoclonal anti-CHI3L1 antibody (FRG) targeting exogenous and or endogenous CHI3L1 effectively inhibits the expression of ACE2 and the uptake of pseudoviruses with the delta or other S protein mutations.

**Figure 3.**
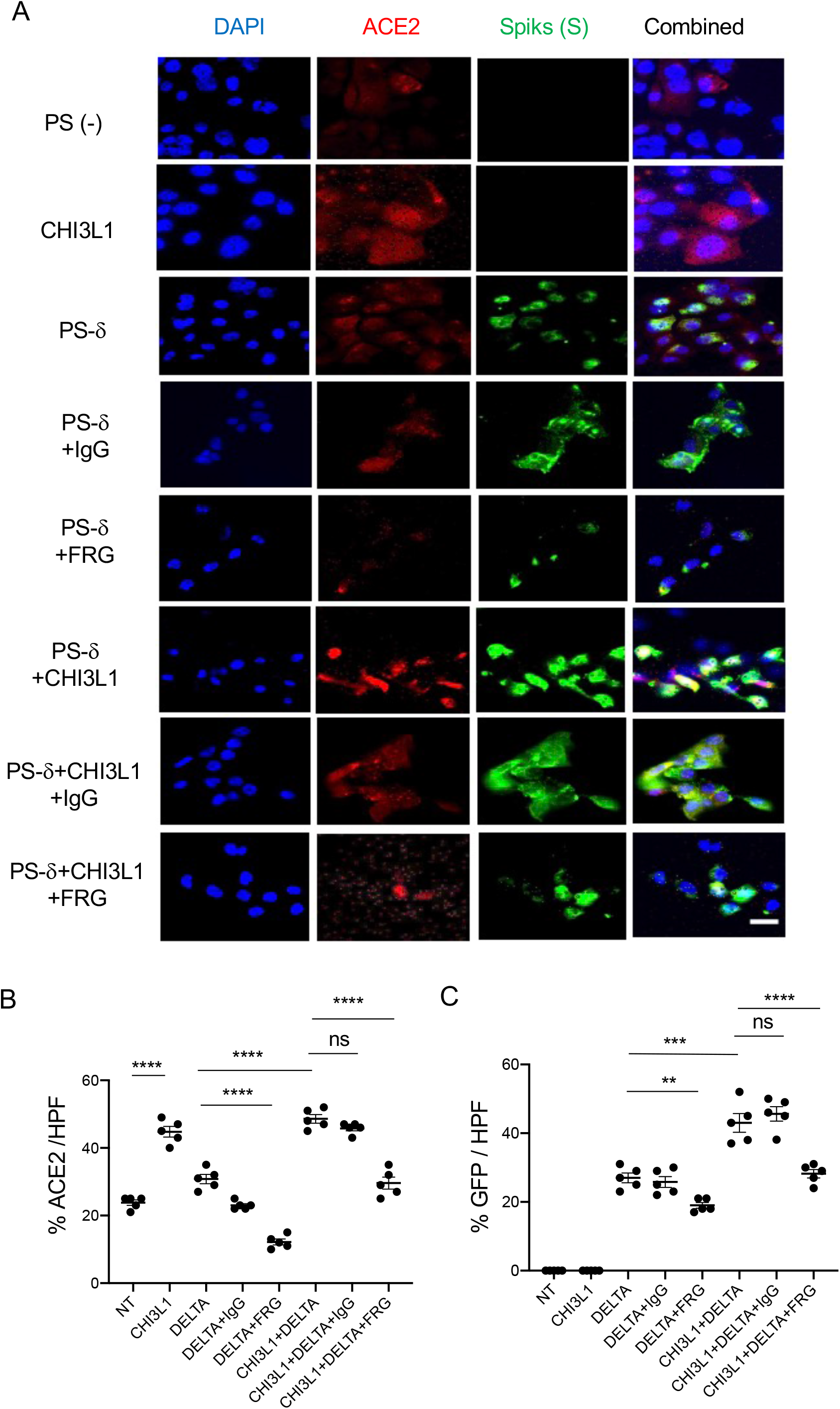
Immunocytochemical evaluation of delta pseudovirus infection of Calu-3 cells. Calu-3 cells were incubated in the presence and or absence of rhCHI3L1 (CHI3L1) in the presence of FRG or its isotype control. (A) Pseudoviruses with delta S proteins (PS-δ) were added and ACE2 and GFP viral infection were evaluated using double labeled immunocytochemistry (ICC). DAPI (blue) was used to evaluate nuclei, red label was used to evaluate ACE2 and the pseudoviruses contained GFP. (B-C) The quantification of ACE2 can be seen in panel (B) and the quantification of GFP is illustrated in panel (C). These evaluations were done using fluorescent microscopy (x20 of original magnification). In these quantifications, 5 randomly selected fields were evaluated. The values in panels B & C are the mean±SEM of the noted 5 evaluations.\*\**P*<0.01; \*\*\**P*<0.001, \*\*\*\**P*<0.0001; ns=not significant (One way ANOVA with multiple comparisons). Scale bar:10µm (applies to all subpanels in Figure 3A).

### Kasugamycin is a small molecule with strong anti-CHI3L1 activity

Recent studies from our lab and others identified that Kasugamycin, an aminoglycoside antibiotic, is a novel small molecule that has a strong anti-Chitinase 1 (CHIT1) activities (27, 28). Since CHIT1 and CHI3L1 share structural homologies as members of 18-glycohydrolase family, we tested whether Kasugamycin (KSM) inhibits CHI3L1 activity similarly to CHIT1. As shown in Figure 4, KSM treatment abrogated CHI3L1 stimulated ERK and AKT activation in Calu-3 cells, suggesting a strong anti-CHI3L1 activity of KSM.

**Figure 4.**
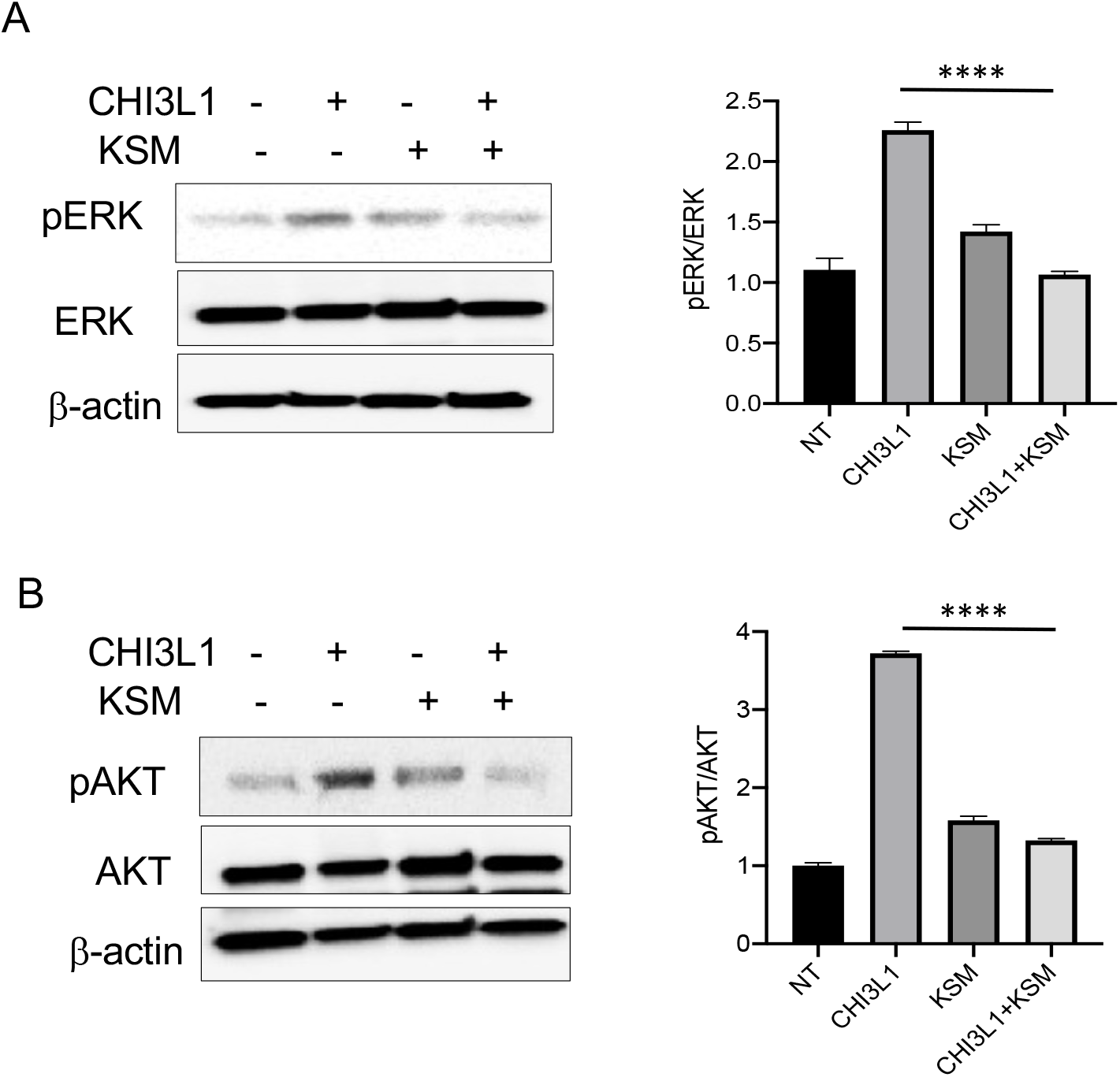
Kasugamycin inhibition of CHI3L1-induced signaling. Calu-3 cells were stimulated with rhCHI3L1 (250ng/ml) or its vehicle control for 2 hours in the presence of Kasugamycin (250ng/ml) and vehicle control (PBS). (A-B) Western blotting and densitometry analysis were then employed to evaluate the levels of activated (phosphorylated, p) and total ERK (A) and AKT(B). The noted figure is representative of a minimum of 3 similar experiments. \*\*\**P*<0.0001 (*t* test).

### Kasugamycin abrogates the CHI3L1-induced increase in epithelial cell uptake of the alpha, beta, and gamma pseudovirus variants

Studies were next undertaken to define the effects of kasugamycin on the uptake of pseudovirus by Calu-3 cells treated with or without rCHI3L1. As seen with the ancestral G614 S protein mutation, treatment of Calu-3 cells with rCHI3L1 augmented ancestral pseudovirus uptake and kasugamycin abrogated these stimulatory effects (Figure 5A). rCHI3L1 had similar stimulatory effects in experiments using pseudoviruses with α, β, or γ S protein mutations (Figure 5, B-D) and these stimulatory effects were markedly decreased by kasugamycin (Figure 5, A-D). When viewed in combination, these studies demonstrate that kasugamycin targeting exogenous and or endogenous CHI3L1 effectively inhibits the uptake of pseudovirus with ancestral, α, β, or γ S protein mutations.

**Figure 5.**
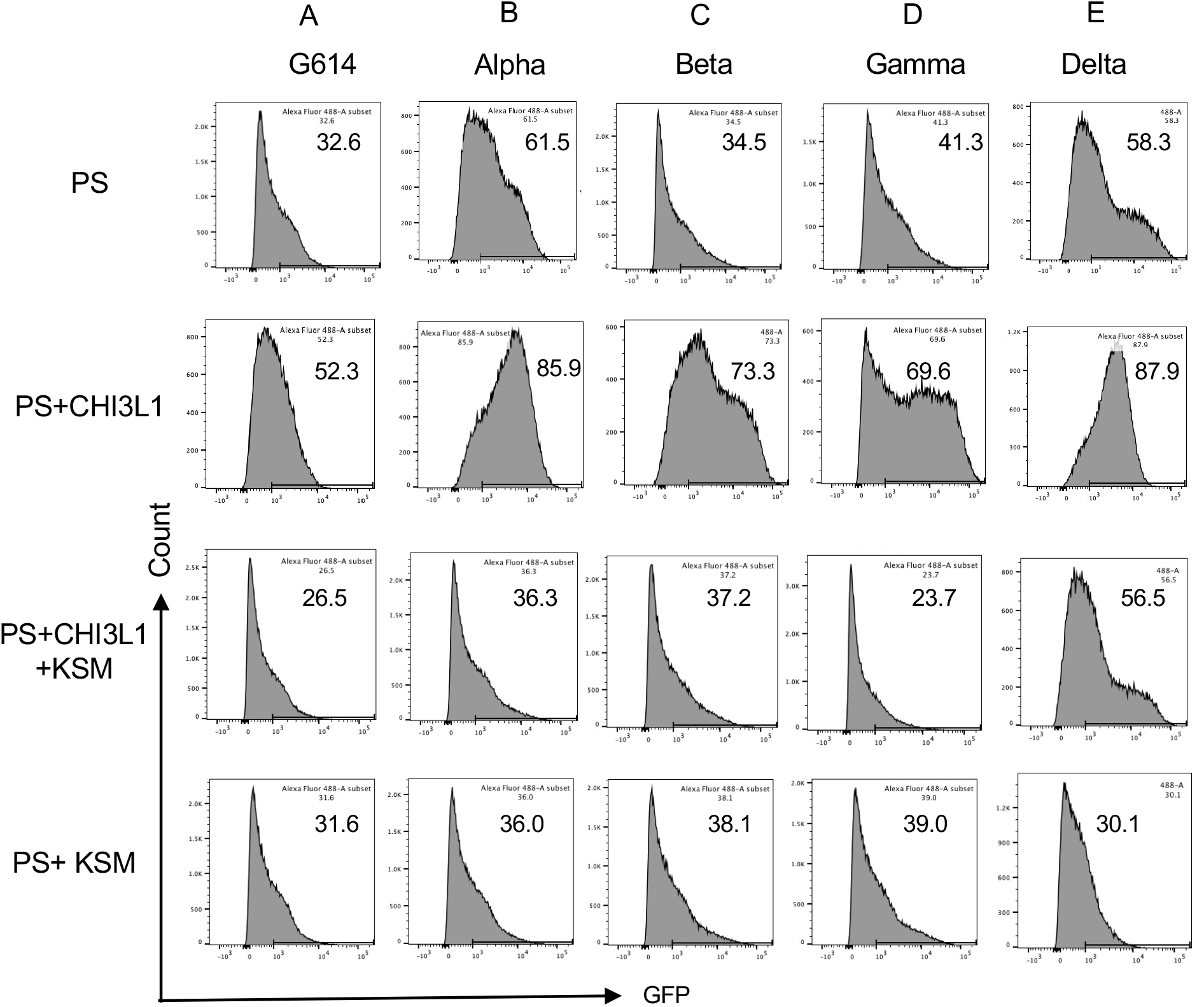
Effects of Kasugamycin on alpha, beta, gamma and delta pseudovirus infection. Calu-3 cells were incubated with rhCHI3L1 (250ng/ml) or vehicle control for 48 hours in the presence of kasugamycin or its vehicle control. Pseudoviruses that contain S proteins with the G614, alpha, beta, gamma or delta mutations were added and GFP was quantitated by FACS analysis. The % of GFP-positive cells was evaluated by flow cytometry. The noted values are representative of a minimum of 3 similar evaluations.

### Kasugamycin abrogates the CHI3L1-induced increase in epithelial cell uptake of delta pseudotyped viral variants

Because of the importance of the delta SC2 viral variant, the ability of kasugamycin to alter the variant’s ability to infect human epithelial cells was also assessed. CHI3L1 was a potent stimulator of the uptake of pseudoviruses that contain the delta S protein mutations (Figure 5E). Kasugamycin abrogated this increase while treatment with the vehicle control did not (Figure 5E). Kasugamycin also diminished pseudovirus uptake by Calu-3 cells even when exogenous CHI3L1 was not administered (Figure 5E). These findings were reinforced by immunocytochemical evaluations. These studies demonstrated that CHI3L1 augmented Calu-3 cell ACE2 accumulation and delta pseudovirus infection (Figure 6). They also demonstrated that FRG abrogated the expression of delta pseudovirus infection at baseline and or after the administration of rCHI3L1 (Figure 6). When viewed in combination, these studies demonstrate that kasugamycin targeting exogenous and or endogenous CHI3L1 effectively inhibits the uptake of pseudovirus with the alpha, beta, gamma or delta S protein mutations.

**Figure 6.**
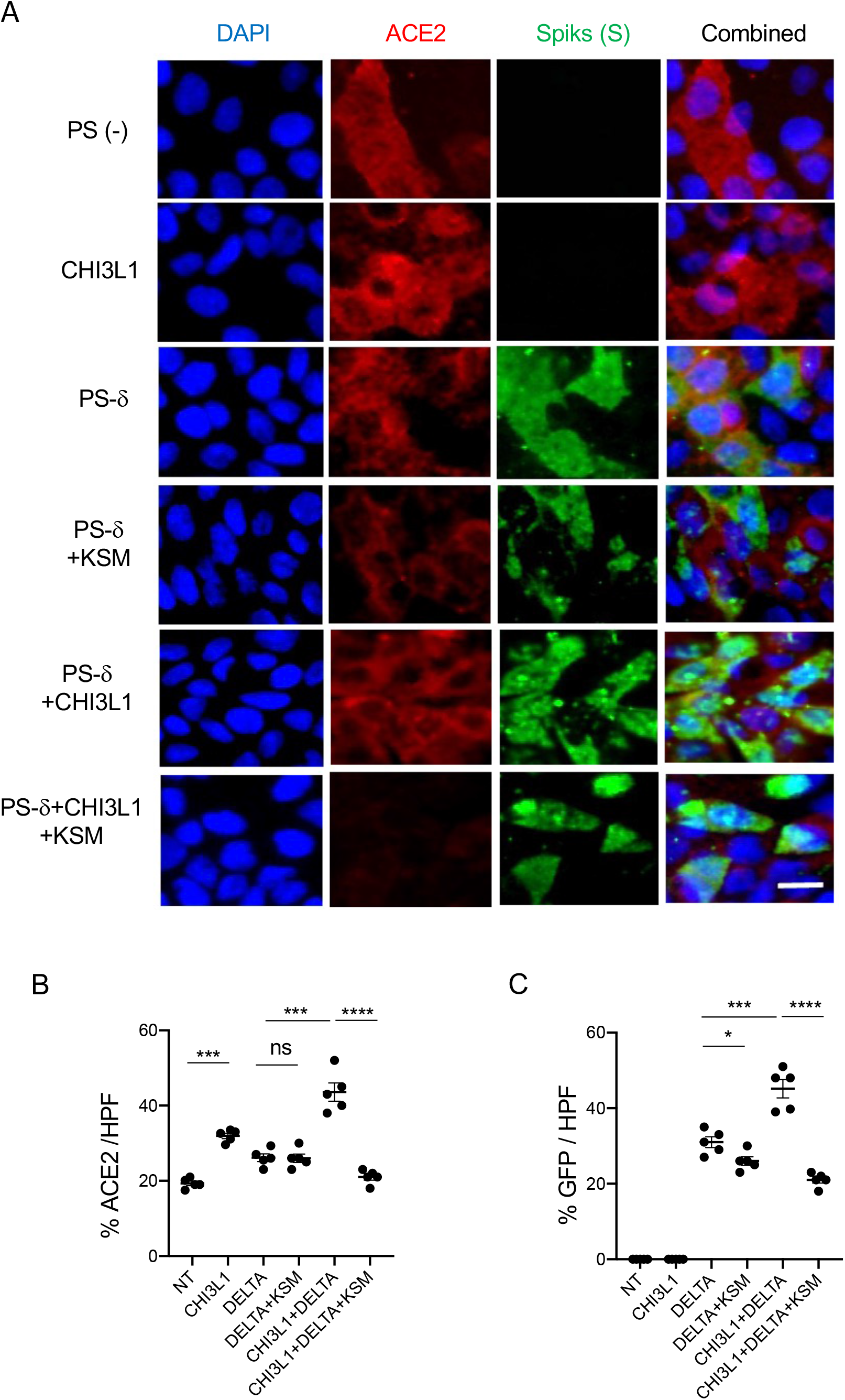
Immunocytotochemical evaluation of delta pseudovirus infection of Calu-3 cells. Calu-3 cells were incubated in the presence and or absence of CHI3L1 (250ng/ml) in the presence or of kasugamycin (250ng/ml) or its vehicle control. (A) Pseudoviruses with delta S proteins were added and ACE2 and GFP viral infection were evaluated using double labeled immunocytochemistry (ICC). DAPI (blue) was used to evaluate nuclei, red label was used to evaluate ACE2 and the pseudoviruses contained GFF. (B-C) The quantification of ACE2 can be seen in panel (B) and the quantification of GFP is illustrated in panel (C). These evaluations were done using fluorescent microscopy (x20 of original magnification). In these quantifications, 5 randomly selected fields were evaluated. The values in panels B & C are the mean±SEM of the noted 5 evaluations. \**P*<0.05, \*\**P*<0.01; \*\*\**P*<0.001; ns=not significant (One way ANOVA with multiple comparisons). Scale bar:10µm (applies to all subpanels in Figure 6A).

### The monoclonal antibody “FRG” and kasugamycin inhibit epithelial uptake of omicron pseudotyped viral variants

Omicron variant has rapidly spread from its first appreciation as a highly mutated variant causing a localized outbreak in South Africa to the most common SC2 variant in the USA and the world (https://covid.cdc.gov/covid-data-tracker/#variant-proportions). Thus, studies were undertaken to determine if the therapies described above that target CHI3L1, ACE2 and SPP in ancestral, alpha, beta, gamma and delta pseudoviruses are also effective in pseudoviruses with omicron S protein mutations. As was seen with the α, β, γ and δ pseudoviruses, CHI3L1 was a potent stimulator of the uptake of Pseudoviruses that contained omicron S proteins (Figure 7). In addition, FRG and kasugamycin both inhibited epithelial cell uptake of pseudoviruses with omicron S protein mutations (Figure 7). These studies demonstrate that antibodies or small molecule inhibitors that target CHI3L1 inhibit epithelial uptake of pseudoviruses with a wide range of S protein mutations including those seen in the omicron SC2 variant.

**Figure 7.**
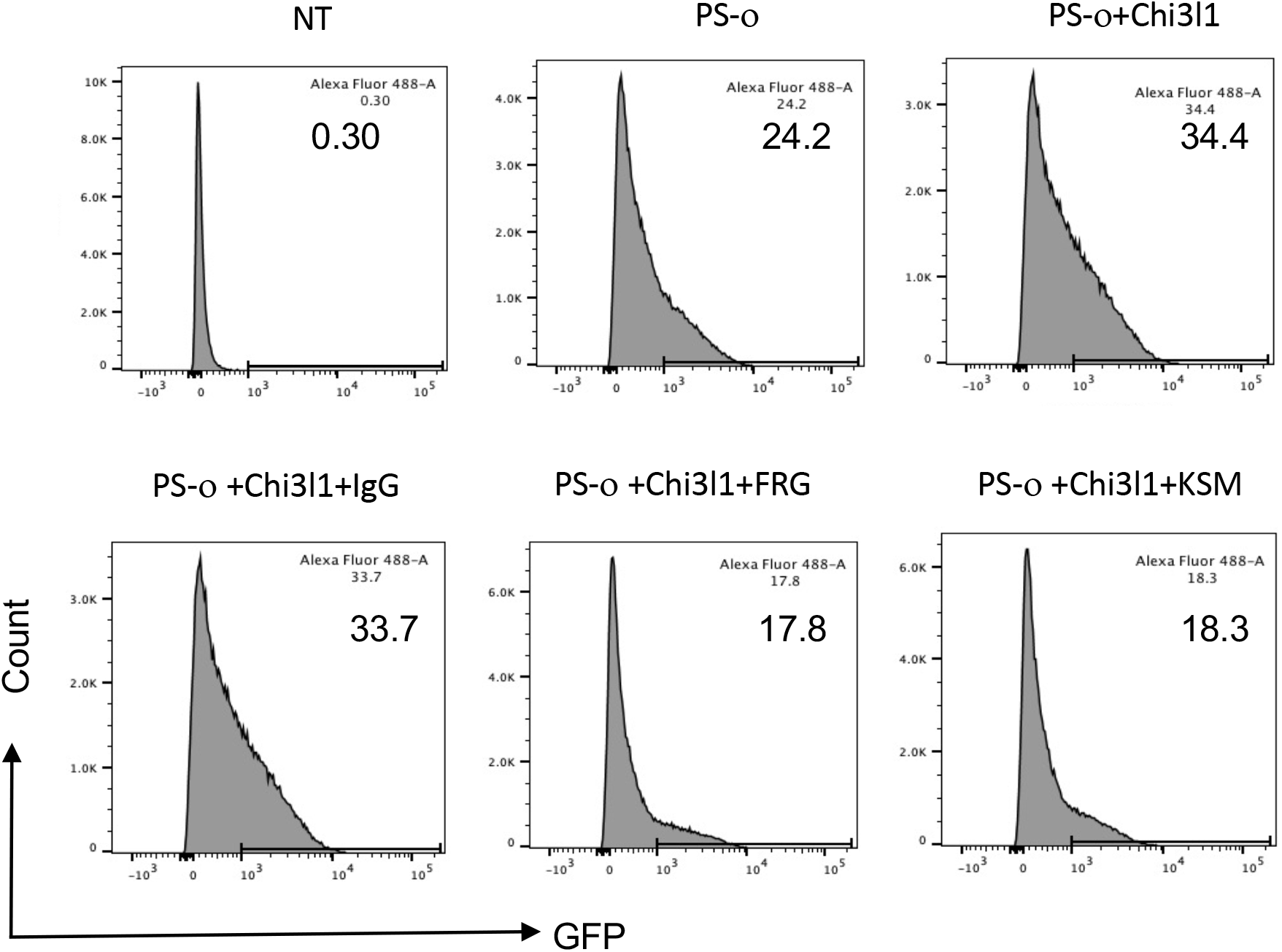
Effects of FRG and Kasugamycin on omicron pseudovirus infection. Calu-3 cells were incubated with rhCHI3L1 (CHI3L1, 250 ng/ml) or vehicle control for 48 hours in the presence or absence of anti-CHI3L1 (FRG) or its isotype control (IgG) or kasugamycin or vehicle control. Pseudoviruses that contain S proteins with the omicron mutations (PS-o) were added and GFP was quantitated by FACS. The % of GFP-positive cells was evaluated by flow cytometry. The noted values are representative of a minimum of 3 similar evaluations.

## DISCUSSION

Coronaviruses are large enveloped single stranded viruses (8). Generally, the rates of nucleotide substitution of RNA viruses are fast and mainly the result of natural selection (29). This high error rate and the subsequent rapidly evolving virus populations can lead to the accumulation of amino acid mutations that affect the transmissibility, cell tropism, pathogenicity and or the responsiveness to vaccinations and or therapies (8, 29). The SC2 VOC are known to manifest enhanced transmissibility and diminished vaccine effectiveness when compared to ancestral controls (30, 31). Their mutations are important causes of viral infection, the cause of new waves of illness and death and drivers of pandemic persistence (30). This can be readily appreciated in the rapid spread of the delta and omicron variants with the latter now accounting for 99.5% of SC2 infections in the USA (https://covid.cdc.gov/covid-data-tracker/#variant-proportions, as of Jan 15, 2021). It can also be seen in delta’s enhanced ability to replicate which drives the viral load up beyond what many other variants can do and outpaces the body’s initial antiviral response (32). The fact that the spectrum of mutations in and characteristics of these variants differs from one another has complicated approaches to vaccination and therapy. In light of the importance of the variants, especially delta and omicron, in COVID 19 studies were undertaken to determine if therapies could be developed by targeting host moieties that help to control many of the major VOC of SC2. In keeping with the importance of ACE2 and SPP in SC2 infection and the impressive ability of CHI3L1 to stimulate these moieties, these studies focused on the relationships between CHI3L1, ACE2 in infections caused by the α, β, γ δ and o variants. They demonstrate that CHI3L1 stimulates the infection caused by these VOC by stimulating the expression and accumulation of ACE2 and SPP. They also demonstrate that antibody-based and small molecule inhibitors of CHI3L1 inhibit the infection of human epithelial cells by these major SC2 VOC including delta and omicron. In combination, they suggest that CHI3L1 is a potential therapeutic target that can be manipulated to prevent or alter the natural history of SC2 infection caused by the current and possible future viral variants that utilize ACE2 and SPP.

The S glycoprotein of SC2 is located on the outer surface of the virion and undergoes cleavage into S1 and S2 subunits. The S1 subunit is further divided into a receptor binding domain (RBD) and an N-terminal domain (NTD) which serve as potential targets for neutralization in response to antisera and or antibodies induced by vaccines (8, 33). Genetic variation in SC2 can have important implications for disease pathogenesis especially if the alterations involve the RBD. In keeping with this concept, the SC2 VOC have impressive mutations in the viral S proteins with alterations in RBD and NTD (34, 35). Three of the VOC have N501Y alterations which augment viral attachment to ACE2 and subsequent host cell infection (8). Omicron has 37 amino acid mutations in its S protein (23). Fifteen of these mutations are in the RBD and nine are in the RBM which is the subdomain of the RBD that directly interacts with ACE2 (23). When viewed in combination, these studies highlight the importance of the viral S proteins and host ACE2 and SPP in the responses induced by SC2 variants. Because our data demonstrates that CHI3L1 is a potent stimulator of ACE2 and SPP they also provide a mechanism by which CHI3L1-based interventions can be effective therapies in all SC2 variants that utilize ACE2 and SPP to mediate viral infection.

Antibodies against the SC2 spike proteins are an evolving and important part of the immune response to SC2 and treatment tool kit against COVID 19. Because the S protein of omicron is heavily mutated, the therapeutic efficacy of vaccine-induced antibodies and commercial monoclonal anti-spike protein antibodies have been characterized. These studies demonstrated that vaccine-induced antibodies can manifest diminished therapeutic efficacy compared to ancestral and other SC2 variants like delta (36, 37). In keeping with these findings, antibodies from Regeneron Inc and Eli Lilly Inc have been noted to manifest diminished potency against omicron while manifesting impressive efficacy against the delta and other variants (36, 37). Our studies demonstrate that the anti-CHI3L1 antibody FRG and kasugamycin, an inhibitor of Chi3l1 decrease the expression of ACE2 and the ability of the ancestral and the alpha, beta, gamma, delta and omicron variants to infect epithelial cells. This led us to hypothesize that FRG and kasugamycin could decrease the infection and spread of all SC2 variants that utilize ACE2 and SPP to elicit cell infection. In keeping with our findings, recent studies have demonstrated that omicron infection requires ACE2 and that omicron binds to ACE2 is more avidly than the binding of delta to ACE2 (38). This supports our contention that interventions that target CHI3L1 can be effective in the treatment of viruses that utilize ACE to infect epithelial cells.

Variants of interest (VOI) are defined as viral variants with specific genetic markers that may alter the transmissibility and or susceptibility of the virus to vaccination or therapeutic interventions when compared to ancestral strains (8). If the features of the variants are subsequently appreciated to exist, the variant is then reclassified as a VOC. As of June 22, 2021, there were 7 VOI including epsilon, zeta, eta, theta, kappa, and lambda. More recently epsilon and Mu have been reclassified as a VOC (8). In all cases ACE2 is presumed to be needed for infection by these viral variants. In keeping with this presumption, we believe that CHI3L1 will also regulate VOI infection, replication and symptom generation by altering ACE2 and SPP. Additional experimentation, however, will be required to formally define the roles of CHI3L1 and effects of CHI3L1 blockade on the effects of these moieties.

Studies from our laboratory and others have demonstrated that CHI3L1 is a critical regulator of inflammation and innate immunity and a stimulator of type 2 immune responses, fibroproliferative repair and angiogenesis (39-46). These studies also demonstrated that CHI3L1 is increased in the circulation of patients that are older than 60 years of age and patients with a variety of comorbid diseases including obesity, cardiovascular disease, kidney disease, diabetes, chronic lung disease and cancer (41, 47-57). In keeping with these findings, we focused recent efforts on the development of CHI3L1-based interventions for these disorders. One of the most effective was a monoclonal antibody raised against amino acid 223-234 of CHI3L1 which is now called FRG. There are a number of reasons to believe that FRG can be an effective therapy in COVID 19. First, as noted by our laboratory (26) and in the studies noted above, it is a potent inhibitor of CHI3L1 stimulation of ACE2 and SPP which decreases the infection of epithelial cells by SC2. In addition, CHI3L1 is a potent stimulator of type 2 immune responses and type 2 and type 1 immune responses counter regulate each other. As a consequence, anti-CHI3L1 augments type 1 immune responses which have potent antiviral properties. Anti-CHI3L1 also inhibits the abnormal fibroproliferative repair responses that are seen in pathologic tissue fibrosis such as that seen in lungs from patients with COVID 19 who require prolonged mechanical ventilation. The present studies add to these insights by highlighting the ability of FRG to inhibit the infection of epithelial cells by the alpha, beta, gamma, delta and omicron SC2 VOC. When viewed in combination, these studies suggest that FRG is a potent therapeutic that can be used to prevent or diminish SC2 infection and or the COVID 19 disease manifestations induced by SC2 and its major variants while augmenting type 1 antiviral responses and controlling tissue fibrosis.

REGEN-COV-2 is a combination of the monoclonal antibodies casirivimab and imdevimab that bind to non-competing epitopes of the RBD of the S protein of SC2 (58). When administered via a subcutaneous route iREGEN-COV2 markedly decreases the risk of hospitalization or death among high-risk persons with COVID 19 (58). Subcutaneous REGEN-COV2 also prevents symptomatic infection in previously uninfected household contracts of infected persons and decreases the duration of the symptoms and the titers of the virus after SC2 infection (58). Because the SC2 VOC have S protein mutations that involve the RBD, one can appreciate the importance of combining two antibodies that target different RBD epitopes to allow these antibodies to neutralize the various VOC including alpha, beta, gamma, delta and epsilon (58-61). Because FRG and casirivimab/imdevimab control SC2 via different mechanisms, it is tempting to speculate that additive or synergistic antiviral and or anti-disease effects including preexposure and postexposure prophylaxis will be seen when FRG and REGEN-COV2 are administered simultaneously. One can also see how the administration of FRG and REGEN-COV2 in combination could protect against the selection of resistant SC2 variants (58).

Kasugamycin was discovered in 1965 in *Streptomyces kasugaensis* and has proven to have antibacterial and antifungal properties (62, 63). Since the 1960s it has been employed as a pesticide to combat agricultural diseases like rice blast fungus and, as a result, has been extensively studied by the Environment Protection Agency (EPA) (64). Most recently Kasugamycin was shown to inhibit influenza and other viral infections (65). Previous studies from our laboratory have added to our understanding of kasugamycin by demonstrating that it is a powerful inhibitor of CHI3l1 induction of ACE2 and SPP that also inhibits type 2 adaptive immune responses and pathologic fibrosis (26, 27). Importantly, the studies in this submission go further by demonstrating that these CHI3L1-based effects of kasugamycin can be seen in the ancestral and alpha, beta, gamma, delta and omicron SC2 VOC. When viewed in combination, these observations suggest that kasugamycin can be used as a prophylactic or therapeutic in COVID 19. This is an interesting concept because kasugamycin can be given via an intravenous or oral route and is known to have minimal toxicity in man (62, 66).

At the onset of the SC2 pandemic there was an urgency to mitigate this new viral illness. Since then significant progress has been made in the treatment of COVID 19 due to intense research efforts that resulted in novel therapeutics and vaccine development at an unprecedented rate (8). The progress that was made, however was diminished by the appearance of SC2 viral variants, particularly delta and omicron. It is now known that SC2 infection results in a disease with two phases. The early phase is characterized by cell infection and viral replication and the latter phase is characterized by a robust host antiviral immune response (8). Current, therapies that are used in the early phase of SC2 infection include antivirals like remdesivir and anti-SC2 monoclonal antibody pairings like bamlanivimab/etesvimab and casirivimab/imdevfimab. When inflammation and a robust immune response have been triggered anti-inflammatories like dexamethasone and immunomodulators are available. The present studies add to our understanding of the therapies for the early phase of SC2 by demonstrating that the inhibition of ChI3L1 with FRG and or Kasugamycin ameliorates the infection induced by the alpha, beta, gamma, delta and omicron SC2 VOC. This raises the exciting possibility that FRG or Kasugamycin, alone or in combination with each other or other SC2 monoclonals can have powerful prophylactic effects and or inhibit viral infection in SC2-exposed individuals. They also demonstrate that FRG and Kasugamycin can directly diminish viral replication and, by decreasing viral load, decrease disease pathology and severity. Additional studies of the importance of CHI3L1 and its roles in infections caused by SC2 variants is warranted.

## MATERIALS AND METHODS

### Cell lines and primary cells in culture

Calu-3 (HTB-55) lung epithelial cells were purchased from American Tissue Type Collection (ATCC) and maintained at 37°C in Dulbecco’s modified eagle medium (DMEM) supplemented with high glucose, L-Glutamine, minimal essential media (MEM) non-essential amino acids, penicillin/streptomycin and 10% fetal bovine serum (FBS) until used.

### Generation of monoclonal antibodies against CHI3L1 (FRG)

The murine monoclonal anti-CHI3L1 antibody (FRG) was generated using peptide antigen (amino acid 223-234 of human CHI3L1) as immunogen through Abmart Inc (Berkeley Heights, NJ). This monoclonal antibody specifically detects both human and mouse CHI3L1 with high affinity (kd 1.1×10^−9^). HEK-293T cells were transfected with the FRG construct using Lipofectamine™ 3000 (Invitrogen, # L3000015). Supernatant was collected for 7 days and the antibody was purified using a protein A column (ThermoFisher scientific, # 89960). Ligand binding affinity and sensitivity were assessed using ELISA techniques.

### Infection of pseudoviruses with S protein mutations

Pseudoviruses with wild type S proteins or the S mutations that are seen in the alpha, beta, gamma, and delta variants were purchased from BPS Bioscience Inc (San Diego. CA). The pseudovirus with omicron S protein was obtained from eEnzyme (Gaithersburg, MD, USA). Pseudoviruses containing S protein mutations of COVID variants used in this study can be seen in Supplementary Table S1. These pseudotyped SARS-CoV-2 virus moieties had a lentiviral core expressing green fluorescent protein (GFP) and the SARS-CoV-2 spike protein but lacked core SC2 sequences. We then compared the ability of pseudotyped virus with mutated S and ancestral S proteins to infect untreated and or treated Calu-3 epithelial cells. A plasmid expressing VSV-G protein instead of the S protein was used to generate a pantropic control lentivirus. SARS-CoV-2 pseudovirus or VSV-G lentivirus were used to spin-infect Calu-3 cells in a 12-well plate (931g for 2 hours at 30°C in the presence of 8 µg/ml polybrene). Flow cytometry analysis of GFP (+) cells was carried out 48 h after infection on a BD LSRII flow cytometer and analyzed with the FlowJo software.

### Immunofluorescence Assay (immunocytochemistry)

Immunofluorescent staining was used to assess cellular integration of pseudoviruses associated with expression of ACE2. Briefly, Calu-3 cells were cultured in 4 well chamber slides (10^6 cell/well) for 24 hr then infected with control and pseudoviruses for 48hrs. Then the cells on the slides were fixed, permeabilized, and treated with blocking buffer then incubated with anti-ACE2 antibody (R&D, AF933) for overnight at 4°C. The photographs of cellular immunofluorescence of GFP (+) pseudovirus and Cy-5 (+) ACE2 expression was taken with fluorescent microscopes

### Western blotting (Immunoblotting)

25 µg lung or cell lysates were subjected to immunoblot analysis using antibodies against phosphorylated (p) ERK (pERK), total ERK(ERK), Phosphorylated(p) AKT (pAKT), total AKT(AKT) (Cell Signaling Tech, MA, USA). These samples were gel fractionated, transferred to membranes, and evaluated as described previously by our laboratory (67).

### Quantification and Statistical analysis

Statistical evaluations were undertaken with GraphPad Prism software. As appropriate, groups were compared with 2-tailed Student’s *t* test or with nonparametric Mann-Whitney U test. Values are expressed as mean±SEM. One-way ANOVA or nonparametric Kruskal-Wallis tests were used for multiple group comparisons. Statistical significance was defined as a level of *P* < 0.05.

## Acknowledgements

This work was supported by National Institute of Health (NIH) grants PO1 HL114501(JAE), and R01 HL115813 (CGL) from NHLBI. This work was also supported by COVID-19 Research Seed Grant from Brown University (CGL).

