## Supplemental Table 1 for "Host Chitinase 3-like-1 is a Universal Therapeutic Target for SARS-CoV-2 Viral Variants in COVID 19"

**Table S1. Pseudoviruses containing S protein mutations of COVID variants used in this study**

| Variant Name | Mutations in S protein | Pseudovirus Source (Cat #) |
|---|---|---|
| Alpha (B.1.1.7 and Q lineages) | Deletions of H69, V70, and Y144; N501Y, A570D, D614G, P681H T716I, S982A, D1118H | BPS BIOSCIENCE Inc. Cat#78112-1 |
| Beta (B.1.351 and descendent lineages) | L18F, D80A, D215G, R246I, K417N, E484K, N501Y, D614G A701V | BPS BIOSCIENCE Inc, Cat #78142-1 |
| Gamma (P.1 and descendent lineages) | L18F, T20N, P26S, D138Y, R190S K417T, E484K, N501Y, D614G H655Y,T1027I | BPS BIOSCIENCE Inc. Cart #78144-1 |
| Delta (B.1.617.2 and AY lineages) | T19R, G142D, 156/157 Deletion, R158G, L452R,T478K, D614G, P681R, D950N | BPS Bioscience Inc. Cat# 78216-1 |
| Omicron (B.1.1.529 and BA lineages) | S371L, G339D, S375F, S373P, K417N, N440K, G446S, S477N, T478K, D614G, E484A, Q498R, H505Y, N501Y, Q493R | eEnzyme.com SCV2-PsV-Omicron |
